# Epitope containing short peptides capture distinct IgG serodynamics that enable DIVA for live-attenuated vaccines

**DOI:** 10.1101/756080

**Authors:** Qinghong Xue, Hongke Xu, Huaidong Liu, Jiaojiao Pan, Jiao Yang, Miao Sun, Yanfei Chen, Wenwen Xu, Xuepeng Cai, Hongwei Ma

**Affiliations:** China Institute of Veterinary Drug Control, Beijing 100081, China; Division of Nanobiomedicine, Suzhou Institute of Nano-Tech and Nano-Bionics, Chinese Academy of Sciences, Suzhou 215123, China

## Abstract

Differentiating infected from vaccinated animals (DIVA) strategies have been central enabling techniques in several successful viral disease elimination programs. However, owing to their long and uncertain development process, no DIVA-compatible vaccines are available for many important diseases. We report herein a new DIVA strategy based on hybrid protein-peptide microarrays which can theoretically work with any vaccine. Leading from our findings from Peste des petits ruminants (PPR), we found 4 epitope containing short peptides (ECSPs) which have distinct IgG serodynamics: anti-ECSP IgGs only exist for 10-60 days post vaccination (dpv), while anti-protein IgGs remained at high levels for >1000 dpv. These data enabled design of a DIVA diagnostic microarray containing 4 ECSPs and 3 proteins, which unlike cELISA and VNT, enables ongoing monitoring of serological differences between vaccinated individuals and individuals exposed to the pathogen. For 50 samples after 60 dpv, 20 animals were detected with positive anti-ECSP IgGs, indicating recent infections in vaccinated goat/sheep herds. These DIVA diagnostic microarrays will almost certainly facilitate eradication programs for (re-)emerging pathogens and zoonoses.

## Introduction

The management of viral diseases in both humans and veterinary medicine involves a series of successive steps including, identification of the causal virus by researchers, treatment by clinicians and veterinarians, public health and agricultural management of active infections (including quarantine and culling), the development and deployment of vaccines to prevent further infections in the population, technologies to enable ongoing surveillance of new viral infections and immunization efficacy, and ultimately the elimination or even global eradication of a particular pathogen. The only two pathogens for which all of these steps have been accomplished are the smallpox and rinderpest viruses ^1, 2^.

However, a notable unintended consequence is that rinderpest eradication may have facilitated the spread of another major viral pathogen, the Peste des petits ruminants virus (PPRV), among sheep and goat herds across Africa and Asia ^3^. Animals exposed to a vaccine against rinderpest are known to become immune to PPRV ^4^, so the cessation of vaccination upon virus eradication suddenly exposed millions of animals to this less deadly but still extremely economically damaging viral disease. Indeed, the most recent global pandemic of PPRV coincided with the vaccination cessation phase of the rinderpest eradication program, a situation that has—based on extensive work by scientists in many national-level programs—led to PPRV’s status as the recognized likeliest candidate for the third successful virus eradication ^5^. Unlike the conspicuously symptomatic smallpox disease and the extremely high lethality of rinderpest, a major challenge for PPR eradication efforts has been the need for ongoing surveillance in herds. In particular the identification of active infections and monitoring of post-vaccination immune status within populations that have already been vaccinated is recognized as the primary remaining barrier to PPRV eradication.

Upon deciding to attempt the elimination of a virus from a particular geographical area, policy managers must select a means to accomplish ongoing surveillance of viral infections in a vaccinated population. Ideally, such a surveillance method would enable monitoring of presence of the virus in a way that distinguishes the serological differences between vaccinated individuals and individuals that have actually been exposed to the pathogen; this would allow for the identification of the precise location where a re-emergence of a pathogen is occurring. To this end, there is a suite of technologies and strategies that are broadly known as “differentiating infected from vaccinated animals (DIVA) ^6^”. A pillar of DIVA strategies employed to date, for example in pseudorabies (PR) ^7^ and foot-and-mouth disease (FMD) elimination programs ^8^, has been the need to develop both an effective DIVA-compatible vaccine (also known as a ‘marker’ or ‘tagged’ vaccine) and a serological test that can differentiate between vaccine-induced antibodies and antibodies against the current field strain of the virus. However, DIVA-compatible vaccines are often not as effective as conventional vaccines, and their development is both time consuming and full of uncertainty ^9^.

For the aforementioned PPRV, the Office International Des Epizooties currently recommends two methods for PPRV diagnosis: a competitive ELISA (cELISA) and a virus neutralization test (VNT) ^10^; unfortunately, both of these are incompatible with a DIVA strategy. Further, there are extensive challenges facing the development and deployment of DIVA-compatible vaccines and corresponding monitoring (S1 Table): 1) taking PR as an example, when a new mutated virus emerges that can escape from the protection provided by the DIVA-compatible vaccine ^11^, the development of a new DIVA-compatible vaccine would not be a timely and would not be an economically wise choice. 2) Taking FMD as an example, there will be detectable anti-NSP (non-structural protein, NSP) IgGs after multiple rounds of vaccination due to residual NSPs in the vaccine, causing a false infection result ^12^. 3) Taking PPR as an example, the negative marker strategy used for DIVA-compatible vaccines is no longer functional for live-attenuated vaccines. To avoid these disadvantageous scenarios, a new strategy is desired.

Peptide microarrays were first developed in the early 1980’s ^13^ after the invention of solid phase synthesis of peptides, a breakthrough that won the Nobel Prize for Chemistry in 1984. After years of exploration, some technical difficulties and high costs have limited the popular application of peptide microarrays to applications such as pharmacological epitope discovery. While working with very low background signal peptide microarray technology ^14, 15^, we recently discovered that epitope containing short peptides (ECSPs) capture different IgG serodynamics than do protein-based assays. In this report, through longitudinal sera analysis with a PPRV-originated peptide microarray, a commercial cELISA, and VNT, we identified 4 dominant ECSPs and confirmed that the distinct IgG serodynamics we initially observed also occur in goats that have been vaccinated with a PPR live-attenuated vaccine. Building from this, we illustrate an entirely new DIVA strategy based on these hybrid ECSP and protein microarrays that, importantly, no longer requires 1) the development of a DIVA-compatible vaccine or 2) the need for negative marker-specific serological monitoring technology. Our work thus opens a large new opportunity for epizoology and zoonoses, raising the possibility that all vaccines can potentially be used in DIVA strategies to facilitate elimination and eradication programs.

## Results and Discussion

Aiming to recapture the different IgG serodynamics detected by protein/virus or short-peptide based assays, we first prepared longitudinal sera by immunizing 9 goats with a live-attenuated vaccine in the laboratory (Fig 1a). The sera were then evaluated using three different analytical platforms, namely our peptide microarray (Microarray-#1), a commercial cELISA, and VNT (Fig 1b-d). Compared to the single (aggregate) index value from a protein based assay ^16^, Microarray-#1 with its 183 peptides would produce 183 separate indices from one sample at a given time point (Fig 1e, left), which is obviously more informative than the other platforms regarding IgG composition in sera. cELISA uses a purified recombinant N protein as the antigen ^17^, and a specific monoclonal antibody, binding to the domain on the amino-terminal half ^18^, as the detection antibody (Fig 1c). The VNT detected antibodies against the F or H proteins that were exposed on the surface of PPRV (Fig 1d) ^19^.

**Fig 1.**
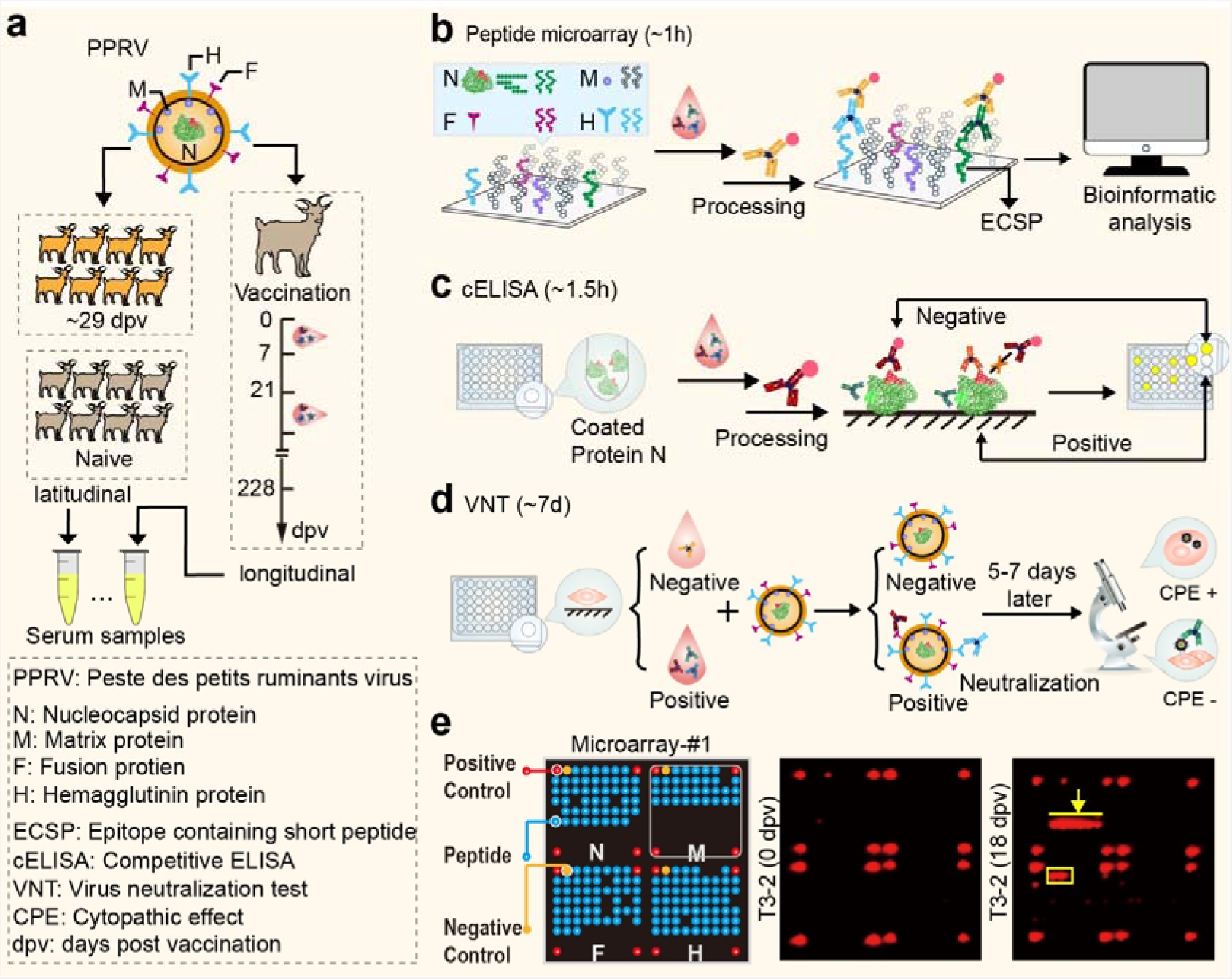
Screening for ECSPs that possess both unique seroconversion dynamics and diagnostic potential. (a) Two categories of serum samples. Longitudinal samples were prepared through immunization of goats with live-attenuated vaccine, and blood samples were collected at specific intervals; latitudinal samples included 92 vaccinated sera at ~29 dpv and 93 negative sera from naive goats without vaccination or infection. (b) Overview of screening with the peptide microarray platform; it is a modified indirect ELISA configured with chemiluminescence, and the whole test takes ~1 hour. Workflow of (c) cELISA (~1.5 hours) and (d) VNT (~7 days). (e) At left, the schematic of Microarray-#1, see S1 Fig for pictures of real objects. It constitutes four subarrays; each subarray contains four positive control points (red, goat IgG), one negative control point (yellow, printing buffer), and 47, 31, 49, and 56 20-mer peptides from the N, M, F, and H proteins of PPRV, respectively (blue). The missing blue bots within each subarray indicate that the corresponding peptides were note able to be synthesized. At middle and right, two representative chemiluminescence images from the peptide microarray with longitudinal samples at 0 dpv (middle) and 18 dpv (right) from the T3-2 goat.

Subject T3-2 was chosen as a representative case based on its relatively higher signal level indicated by the results of Microarray-#1 (Fig 2a). Compared to the negative control serum sampled at 0 dpv (Fig 1e, middle), two groups of peptides, N45-N50 from N protein (Fig 1e, right, yellow arrow) and F9-F11 from F protein (Fig 1e, right, yellow box), were observed over time during the immune response. A large scale analysis with a clustered heatmap (Fig 2b) clearly demonstrates that these peptides contain linear B-cell epitopes. No such peptides were found for the M or H proteins. All 8 other goats showed the same trend, with only minor dissenting features due to individual immune differences ^20^ (S2-4 Fig), so we henceforth focused on N45-N50 and F9-F11 as ECSPs.

**Fig 2.**
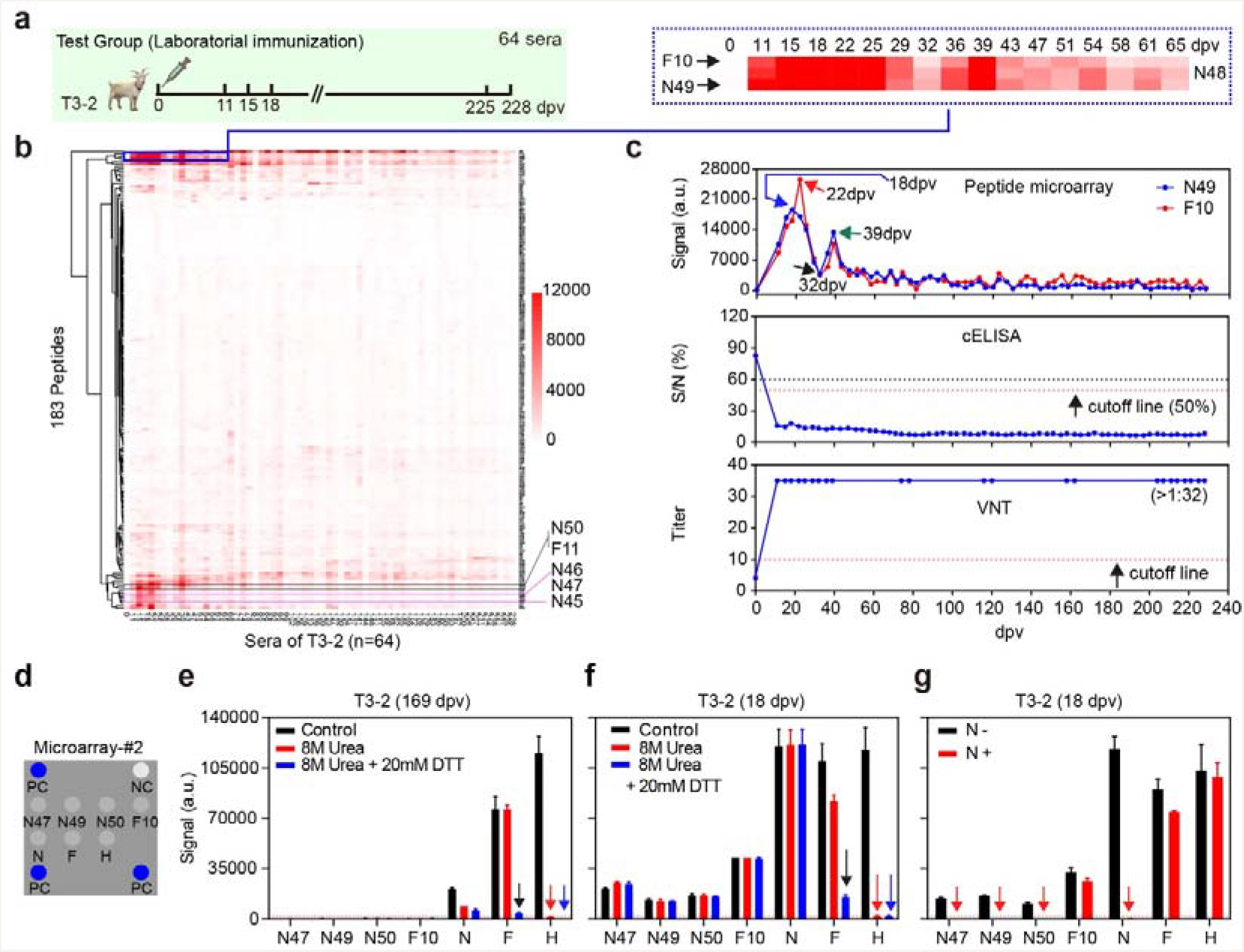
Distinct dynamics and epitope type changes of IgGs. (a) 64 longitudinal sera collected from a representative goat (T3-2). (b) The clustered heat-map of the 64 longitudinal sera from T3-2 against 183 peptides (Microarray-#1). (c) Comparison of the seroconversion dynamics from 64 longitudinal sera of T3-2 detected by ECSPs (N49 and F10), cELISA kit, and VNT, respectively. (d) The layout of Microarray-#2: PC is the goat IgG and NC is the printing buffer as, respectively, the positive and negative controls for horseradish peroxidase (HRP) conjugated antibody interaction. Results of serum (e) T3-2 (169 dpv) and (f) T3-2 (18 dpv) incubated with Microarray-#2 after different denaturing treatments. (g) Blocking experiment with N protein incubated with T3-2 (18 dpv). Treatment with 20 mM DTT alone showed no signal change on any of the ECSPs and proteins (data not shown).

Previous studies have shown that the PPR live-attenuated vaccine can induce lifelong immunity ^21^. Our results showed that the titers of neutralization antibodies detected by VNT and the non-neutralization IgGs detected by cELISA were maintained at constantly high levels from 11 dpv to 228 dpv (Fig 2c, middle and bottom). However, the IgG serodynamics detected by ECSPs, such as N49 or F10, were dramatically different from cELISA and VNT: with ECSP we noted a bimodal curve with two peaks at the early stage (Fig 2c, top). Specifically, the signal for the anti-N49 IgGs was undetectable at 0 dpv and reached an initial peak at 18 dpv (blue arrow, signal=18,524), then declined to a low but detectable level at 32 dpv (black arrow, signal=3,800), before reaching a second but lower peak at 39 dpv (green arrow, signal=13,372), and finally declining dramatically to below the detection limit by around 60 dpv. The activity of anti-F10 IgGs was similar to anti-N49 IgGs, but reached its first peak at 22 dpv, 4 days later than the anti-N49 IgGs.

Given that the results from Microarray-#1, cELISA (protein N), and VNT (mainly proteins F and H at the surface of PPRV) cannot be compared quantitatively, we printed 4 ECSPs and 3 proteins collectively to form Microarray-#2 (Fig 2d and 3a) for further tests. To explore the position of the ECSPs on the proteins, we performed blocking experiments (S5c Fig) and the results not only confirmed their specificity, but also supported that the three N-protein ECSPs and F10 are located on the surface of protein N and F. For example, free ECSP N47 blocked the signal of N47 in Microarray-#2, but did not block other ECSPs or proteins (S5d Fig). Additionally, since free protein N was able to block both protein N and the ECSPs which originated from protein N in Microarray-#2, we concluded that the C terminus of protein N is rich in linear epitopes, similar to protein N of the measles virus (MeV); both viruses belong to the same Morbillivirus genus ^22^ (Fig 2g and S5d Fig).

**Fig 3.**
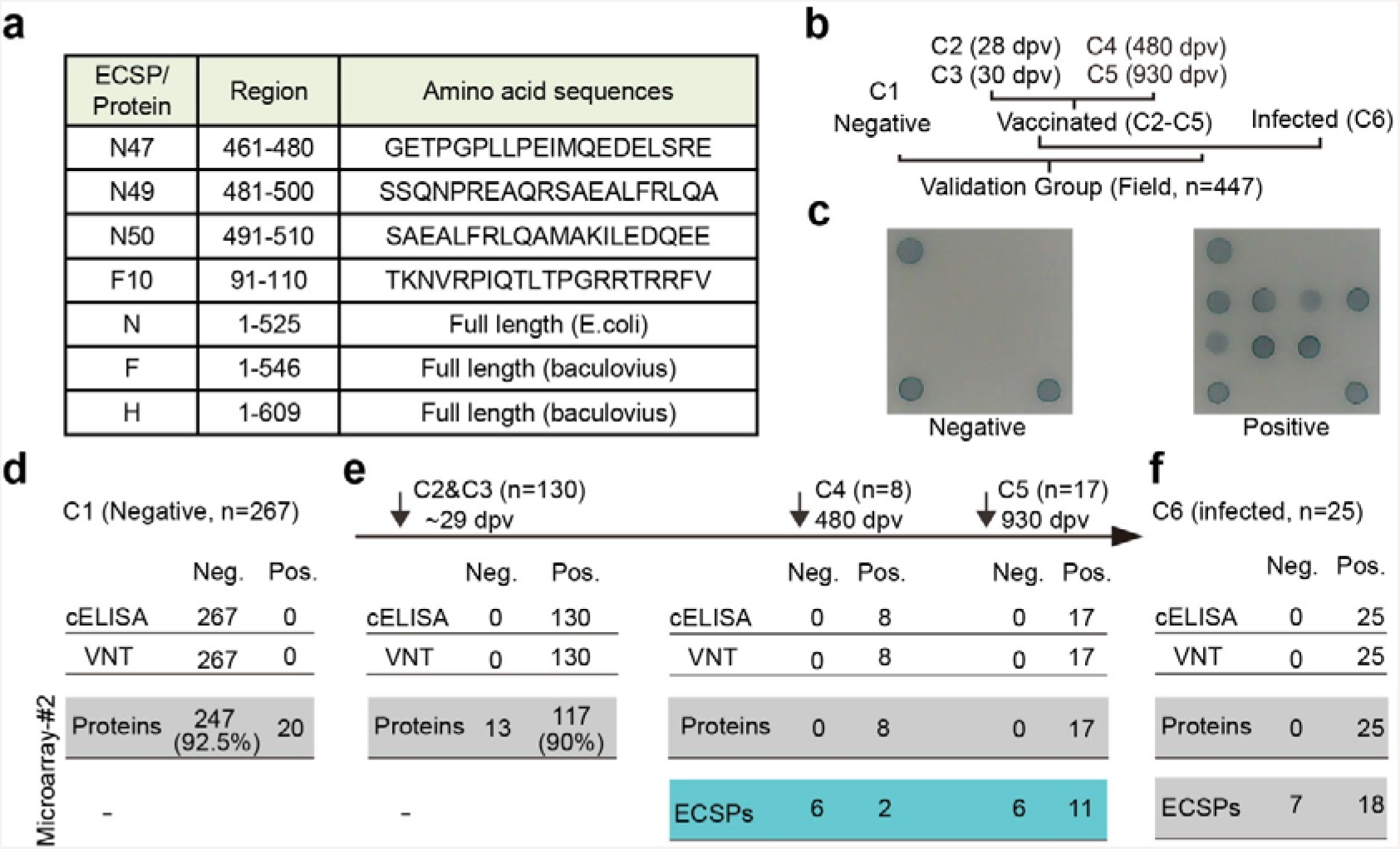
Validation of distinct IgG serodynamics, using latitudinal sera, which have implications for vaccine titer evaluation and DIVA. For individual ECSPs or proteins of Microarray-#2, the result was positive if its S/P was larger than the cutoff, see S10 Fig for details. For the ECSPs combination (i.e. N47+N49+N50+F10), the result was positive if any ECSP was positive. For microarray-#2, the result was positive if any two proteins were positive (mainly for titer evaluation) or if any ECSP was positive (mainly for DIVA). (a) Selected ECSPs and proteins. (b) Six groups of sera (C1-C6) were collected to test the Microarray-#2. The positive/negative statuses of these latitudinal samples were confirmed by both cELISA and VNT. (c) Left and right are representative images of negative and positive serum. Results of sera against microarray-#2: (d) C1, the 267 negative sera; (e) Left: 130 positive sera (C2 and C3 combined); Middle: C4, 8 positive sera at 480 dpv; Right: C5, 17 positive sera at 930 dpv; (f) C6, 25 positive sera due to virus infection.

The different IgG serodynamics detected by ECSPs and cELISA/VNT may be attributed to the short and long durations of IgGs that recognize linear and conformational epitopes, respectively. To probe the nature of interactions between ECSPs/proteins and IgGs at different immunization stages, prior to incubation with serum, we used the disulfide-reducing agent Dithiothreitol (DTT) and protein denaturant urea to disrupt conformations of printed proteins in Microarray-#2 (S5a-d Fig).

First, using intact proteins as control, without denaturing treatment (the black columns, Fig 2e-f) gave a quantitative comparison of various IgG levels. A later stage serum T3-2 (169 dpv) lacked anti-ECSP IgGs but contained different levels of anti-protein IgGs (Fig 2e). Signals due to anti-H, anti-F, and anti-N IgGs were ~115,300, ~76,600 and ~20,950, respectively. An early stage serum T3-2 (18 dpv) contained IgGs for both ECSPs and proteins (Fig 2f). Signals for anti-ECSPs ranged in intensity from 13,000 to 42,500; we also measured the signals for anti-H (~118,000), anti-F (~109,900), and anti-N IgGs (~120,200).

These quantitative data implied that proteins N, F, and H can out-compete ECSPs. At the early stage, proteins induced 3-10 times more IgG production than ECSPs (assuming all IgGs have the same affinity such that the detected signal depends solely on concentration). At the later stage, proteins induced long-lived plasma cells (LLPC) for a continuous supply of IgGs, which is the key to obtaining lifelong immunity. While the anti-N IgG level dropped dramatically (from ~120,200 to ~20,950), the anti-H IgG level was almost constant (~115,300 vs. ~118,000) and anti-F IgG level dropped slightly from ~109,900 to ~76,600, implying the immune system was able to selectively focus on the production of functional (i.e., neutralization) IgGs ^16^.

Second, printed proteins denatured by combined urea and DTT (hereafter urea+DTT) treatment were used to infer whether the epitope type of IgG changed during the course of the immune response (the blue columns, Fig 2e-f). For a later stage serum T3-2 (169 dpv), the urea+DTT treatment caused a significant signal loss for protein N (from ~20,950 to ~5900), H (~115,300 to 125), and F (~76,600 to ~4,300) (Fig 2e). Such sensitivity to protein conformational changes can be explained by assuming there were only IgGs that recognize conformational epitopes on protein N, H, and F, (or perhaps other IgGs were present but below the detection level). In contrast, for an early stage serum T3-2 (18 dpv) (Fig 2f), we found that none of the four ECSPs exhibited any obvious signal change after urea+DTT treatments, which agreed with the fact that short peptides (20-mer in this paper) have no conformation per se and are thus apparently insensitive to urea+DTT treatments. Three proteins, however, behaved differently: protein N and F had both IgGs for linear and conformational epitopes; protein H had IgGs only for conformational epitopes.

For protein N, almost no signal loss was observed after urea+DTT treatments (Fig 2f). This finding suggested that either the conformational structure of the N protein was not disrupted completely or there were abundant IgGs which can recognize the denatured N protein (i.e., linear epitopes) to compensate for the loss of signal from IgGs which are sensitive to conformation. The later scenario is more likely, given that the primary structure of protein N contains only one cysteine in its full length, and considering that this serum sample represents the peak of the ECSPs’ IgG serodynamics, thus likely possessing abundant IgGs which can recognize both linear and conformational epitopes.

For protein F, the signal dropped from 109,918 to 15,312 with the urea+DTT treatments (Fig 2f). This significant remaining signal indicated the existence of IgGs that recognize linear epitopes, such as F10. For protein H, a >98% loss of signal from the H protein after urea+DTT treatments suggested that the H protein probably has no linear epitopes (Fig 2f, S5d Fig).

Third, printed proteins denatured by urea treatment alone indicated the IgG epitope type changes for the three proteins were different (the red columns, Fig 2e-f). Protein N and H showed no differences in signal loss for either urea or urea+DTT treatments (red and blue columns, Fig 2e-f). Protein F, however, responded differently to urea and urea+DTT treatments: the signal dropped from ~109,900 to ~82,100 after urea treatment and dropped to ~15,300 with the urea+DTT treatments (Fig 2f). We therefore propose that at the early stage there were three groups of IgGs (S5b Fig): one is sensitive to conformation and can be disrupted by urea alone (signal contribution 25%), one is sensitive to conformation and can be disrupted by the urea+DTT treatment (signal contribution 86%), the final one is for linear epitopes (signal contribution 14%). For the later stage serum (Fig 2e), the urea+DTT treatment led almost complete signal loss, indicating there were IgGs for conformations but not for linear epitopes, while the treatment of urea alone showed no signal loss, indicating there were no IgGs for conformations sensitive urea alone.

This conformational sensitivity of IgGs to the F and H proteins implies that these IgGs are very likely responsible for the neutralization function which we observed in the VNT results. These results are in agreement with the current model of IgG development as elicited by immunization ^16, 23, 24^: at the early stage, immunization elicited IgGs can recognize both linear (ECSPs and proteins) and conformational epitopes (proteins). As the immune system continues to interact with antigen (vaccine), the host is able to, via an unknown mechanism, concentrate its resource to produce a few but dominant IgGs that are functional-neutralization IgGs for conformational epitopes present at the surface of a virus. We should therefore be able to both detect IgG epitope type changes and to monitor different IgG serodynamics occurring between ESCPs and proteins over time during an immune response.

Due to the limited number of experimental goats, the longitudinal sera group only had a small number of samples at each time point (3 samples for the T3 group). Attempting to extend our findings about IgG serodynamics, we tested more latitudinal field sera at several time points of their serodynamic curves, namely the C1-C6 groups (Fig 3b), which also served as the verification/validation groups for Microarray-#2. Furthermore, 3,3’,5,5’-tetramethylbenzidine (TMB) was used to replace the chemiluminescence substrate, so the microarray was more affordable for field use. A testing procedure was established with positive and negative control sera included in the test, and the signal of individual ECSPs or proteins was further converted to the ratio of the sample to the positive control (S/P). The cutoff S/P for each ECSP or protein was determined through receiver-operating characteristic (ROC) curve analysis based on 267 negative sera (C1) and 130 vaccinated sera at ~29 dpv (S7 Fig).

According to the aforementioned IgG serodynamics, C1-C3 (early stage sera, < 60 dpv) should have the same result pattern from both ECSPs and proteins. 267 negative sera (C1) were tested for their relative specificity. The relative specificity for the ECSP combination was 90.6% (S8a Fig), but reached 92.5% for the protein combination (Fig 3d), both meeting the industrial standard of specificity, i.e., >90%. For the 20 samples with discrepancy (“false positive”), 10 of them were in the gray-zone (near the cutoff line). 10 of them, however, were confirmed positive by Microarray-#1 (S8d Fig).

For the C2 and C3 groups (~29 dpv), although ~1 week behind the peak of the IgG serodynamic curves of anti-ECSPs, we still obtained a relatively high sensitivity of 90% for ECSPs (we believe the sensitivity may be higher than 90% with samples ~20 dpv, S8a Fig). The relative sensitivity using the protein combination was also 90% (Fig 3e, left). For the 13 samples with discrepancies (“false negative”), all were confirmed negative by Microarray-#1 (S8e Fig). We attributed this discrepancy to immune differences, and are examining this in an ongoing project. Together (C1-C3), we confirmed that the behavior of both ECSPs and proteins in Microarray-#2 agreed with the IgG serodynamic trends. Thus, Microarray-#2 is suitable for titer evaluation of the PPR live-attenuated vaccine.

According to the aforementioned IgG serodynamics, C4-C5 (later stage sera, > 60 dpv) would be expected to have different results between ECSPs and proteins. Thus, we present the results of Microarray-#2 for these groups in the following format: (proteins, ECSPs). For C4 (480 dpv) and C5 (930 dpv), one anticipated (+, −) based on the IgG serodynamic curves, but 2/8 and 11/17 sera for C4 and C5, respectively, showed (+, +) (Fig 3e, right). These 13 serum samples were confirmed by Microarray-#1 (S8f Fig), implying these goats were infected by PPRV within the last 60 days before sampling. The agreement of both Microarrays (−#1 and −#2) suggested that ECSPs captured newly elicited IgGs while the proteins capture IgGs secreted by both short-lived plasma cells (SLPC) and LLPC. Thus, we hereafter refer to Microarray-#2 as the DIVA microarray. For the 25 sera from goats infected by PPRV (as confirmed by both cELISA and VNT) (C6), we did not know the exact date of infection. 7/25 were (+, +), indicating recent infection (< 60 days post infection (dpi)); 18/25 were (+, −), indicating an infection time point > 60 dpi (Fig 3f). Note that this timing information is provided by the DIVA microarray but not cELISA or VNT.

To further examine the diagnostic and DIVA potential of Microarray-#2 with larger sample sizes, and inspired by our results for the C4-C6 groups, we designed a general program (Fig 4a) that realizes DIVA surveillance for both a well-managed herd and a poorly-managed herd/wildlife. For the well-managed herd, one knows the vaccination date, so blood sampling started from Tbx (x=1, b stands for before T0). The results obtained as (proteins, ECSPs) for sera against DIVA microarray should follow the curves in Fig 4b. For the period of 0-60 dpv, the result could also be used to evaluate vaccination titer. The interval between two Ta dates (a stands for after T0) could be adjusted as needed from 3-20 days. After 80 dpv for example, a (+, +) or (+, −) result would indicate, respectively, the presence or absence of infection (Fig 4b).

**Fig 4.**
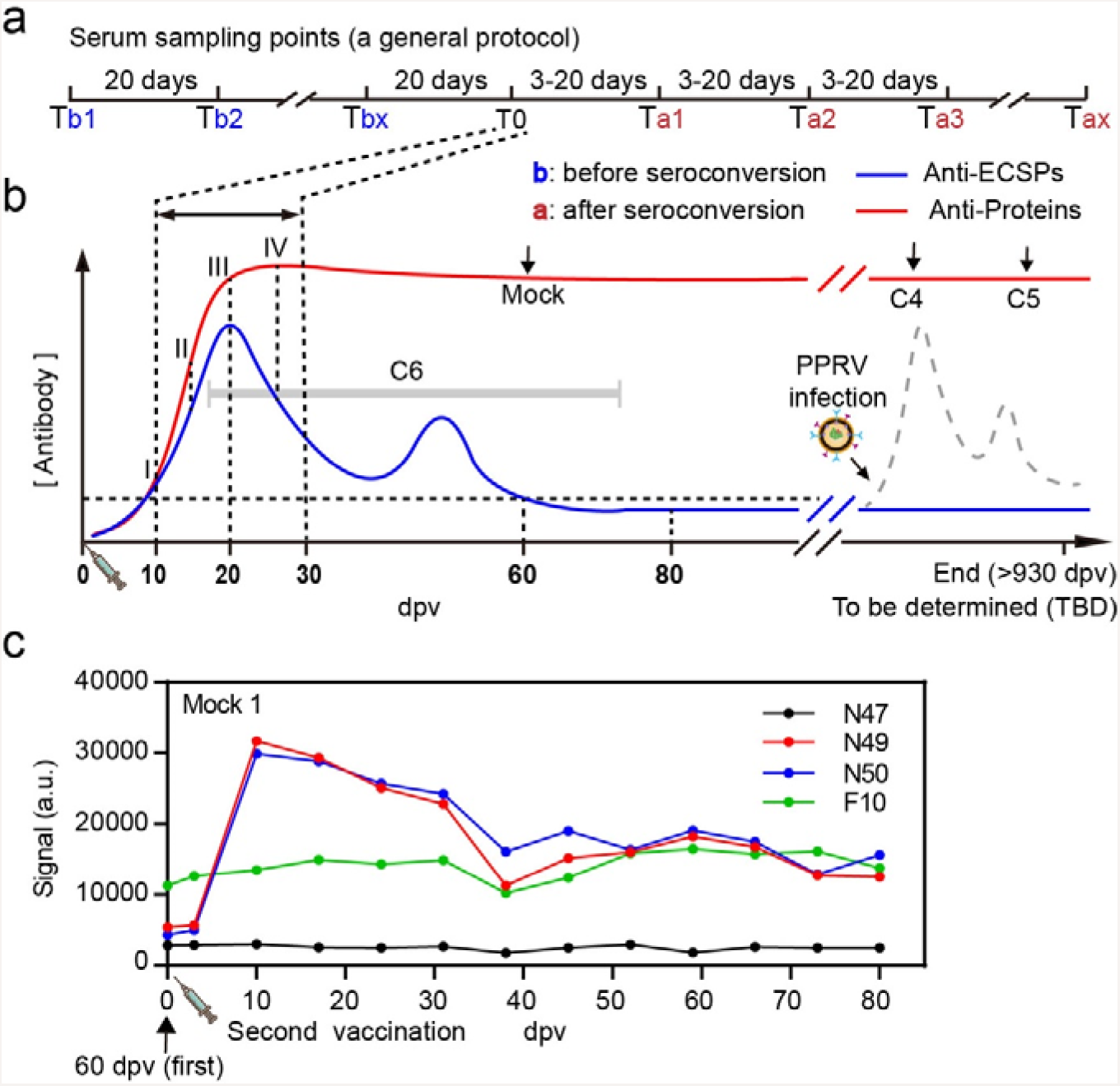
A general protocol for multiple purposes: vaccine titer evaluation, DIVA diagnosis and surveillance as suitable application scenarios. (a) A proposed serum sampling schedule, (b) Serum sampling points matched with IgG serodynamics, (c) Using PPR live-attenuated vaccine to mimic the virus infection, the spiked ECSPs response clearly indicated “infection”.

For a poorly-managed herd, if the vaccination date was unknown (or not vaccinated), represented hypothetically as the gray band of C6 in Fig 4b, a randomly picked date could be used as Tb1. The T0 could be anywhere from 10 to 28 dpv, partially affected by the individual immune status ^20^. We recommended a 3–5 day interval between T0 and Ta1 and Tax (x=60/interval) so a detailed IgG serodynamic curve could be obtained, which would be informative for deducing the position T0.

As a mimic of virus infection, a second immunization (with 50 doses) was carried out on goats at 60 dpv from the first immunization. Although given as a 50-fold increase in dosage, the anti-protein IgG level remained the same (signal = ~120,000). ECSPs N49 and N50 peaked at 10 dpv of the second immunization, and waned monotonically. The signal did not reach zero, even at 80 dpv, which was attributed to the 50-fold increase in dosage (see S10 Fig for another 2 subjects). Since the live-attenuated vaccine is most close to the situation of wild virus infection, this DIVA microarray strategy should in theory work for any vaccine (S2 Table): the protein section detects long lasting IgGs (same function as cELISA and VNT), and the ECSP section determines the stage of vaccination/infection (the new function vital for DIVA). When these two sections are combined, this microarray realizes DIVA. Our ongoing efforts are to translate this DIAV microarray strategy to fight emerging infections and zoonoses ^25–27^. As ever more vaccines are developed to prevent human diseases, e.g., human papillomavirus vaccines for cervical cancer ^28^ and enterovirus 71 vaccines for hand, foot, and mouth disease (HFMD) ^29^, there is an increasing demand for a DIV solution for humans. We noticed that the IgG serodynamics of most human vaccines have been developed based on data from protein/virus based assays ^30–32^. If short peptides share a similar trend in human vaccines, i.e., they wane monotonically, then obviously a similar DIV strategy could be constructed based on our peptide microarray technology, which would be useful in preventing the transmission of the field strain of a virus. This work also implies that applications of these peptide microarrays should provide access to a new world of vaccines and diagnostics for viral diseases.

## Materials and Methods

### Ethics statement

The protocol of animal study was approved by the Committee on the Ethics of Animal Experiments of the China Institute of Veterinary Drug Control (Permit Number: [2016]00114). The study was conducted following the Guide for the Care and Use of Animals in Research of the People's Republic of China.

### Vaccine and vaccination

PPR vaccine is produced by Tiankang Co., Ltd and is a live vaccine based on Nigeria 75/1 attenuated strain (lineage II). The virus titer was 3.2 log10 TCID50/dose of 1 mL. PPR-capripox live vaccine is produced by Sinopharm Yangzhou VAC Biological Engineering Co., Ltd, The PPR virus titer was 3.3 log10 TCID50/dose of 1 mL, and the POX virus titer was 3.6 log10 TCID50/dose of 1 mL. To evaluate the IgG serodynamics elicited by the vaccine, three healthy and susceptible goats (T1) were immunized with PPR vaccine through 1-mL subcutaneous injection and sampled during 0-21 dpv; three goats (T2) were also immunized with PPR vaccine and sampled during 0-44 dpv; three goats (T3) were immunized with PPR-capripox vaccine through 1-mL subcutaneous injection and sampled during 0-228 dpv;

### Virus and challenge

To explore the antibody titer variation of the challenged goat, a second vaccination with live-attenuated vaccine (single vaccine) that mimic the challenge was carried out on the first vaccinated goat (PPR-capripox vaccine) at 50-goat dose was administrated to vaccinate three corresponding goats.

### Peptide library

The proteins of N (CAA52454.1), M (CAJ01698.1), F (CAJ01699.1) and H (CAJ01700.1) from PPR vaccine strain Nigeria 75/1 (NCBI: X74443.2) were employed to analyze the amino acid (aa) sequences. 20-mer peptides with an overlap of 10 aa residues covering the entire protein were chemically synthesized by GL Biochem (shanghai, China), which finally yielded 183 peptides in total, including 47 peptides from N (N1-N52 while N27, N32, N34, N51 and N52 were failed to be synthesized), 31 from M (M1-M33 while M6 and M14 were failed to be synthesized), 49 from F (F1-F54 while F12, F30, F32, F39 and F50 were failed to be synthesized) and 56 from H (H1-H60 while H4, H5, H33 and H46 were failed to be synthesized), respectively.

### Microarrays

Microarray-#1 (peptide microarrays) with whole panel of 183 peptides were fabricated as previously reported (33). Briefly, ~0.6 nL of each peptide with concentration of 0.1 mg/mL were printed onto the activated nano-membranes by contact spotter Smart 48 (Capital Bio, Beijing, China) to form 9×9×4 microarrays (Fig 1e, left). In each sub-array there are four positive controls printed with goat IgG at the concentration of 50 μg/mL and one negative control with printing buffer.

Microarray-#2, including four ECSPs (N47, N49, N50 and F10), three proteins (N, F and H) and goat IgG (i.e., three positive control points) were printed in 4×4 array by non-contact spotter sciFLEXARRAYER S1 (Scienion, Berlin, Germany) (Fig 2d), with the concentration of 0.1 mg/mL for each peptide or protein and 50 μg/mL for goat IgG. N was purchased from Suzhou Epitope Biotech Ltd. Co., China. Protein F and H were purchased from Novo Biotech, China.

### Serum screening with microarrays (Test group)

All the longitudinal samples and part of latitudinal samples including 92 vaccinated sera at ~29 dpv and 93 negative sera from naive goats were first screened using Microarray-#1 as previously described with minor modification ^33^. Serum was first diluted 1:50 with serum-dilution buffer (1% bovine serum albumin,1% Casein, 0.5% Sucrose, 0.2% Polyvinylpyrrolidone, 0.5% Tween20 in 0.01M Phosphate Buffered Saline, pH 7.4) and 200 μL was added into each microarray, incubated for 30 min on a shaker (150 rpm, 22°C). Microarray incubated with serum-dilution buffer was conducted as negative control. The microarray was then rinsed for 3 times with washing buffer and incubated with 200 μL of Horseradish peroxidase (HRP) conjugated Rabbit anti-Goat IgG (Sigma-Aldrich) diluted 1:20000 in Peroxidase Conjugate Stabilizer/Diluent (Thermo Scientific) for another 30 min on a shaker (150 rpm, 22°C), followed by the same washing steps as described above. 25 μL of chemiluminescence substrate (Thermo scientific) was added onto the microarray and the Images were taken at a wavelength of 635 nm using Clear 4 imaging system (Suzhou Epitope, China). The images were analyzed with Matlab. The signal of any peptide dot was defined as signal readout of dot minus signal readout of background.

### cELISA

cELISA (IDvet) was performed according to the manufacture’s instruction. Briefly, 50 μL volumes of samples diluted 1:2 were added into the 96-well plate, where positive and negative controls were in duplicate respectively. The plate then was incubated for 45 min ± 4 min at 37°C (±3°C) and followed by 3 times of wash with wash buffer. 100 μL of conjugate (1×) was added into the wells and incubated for another 30min±3min at 21°C (± 5°C) and followed by 3 times of wash again. 100 μL of substrate was added into the wells and incubated for 15 min ± 2 min in dark at 21°C (± 5°C) and followed by the addition of 100 μL of stop solution. Finally, the absorbance of each well was read at 450 nm using microplate reader.

### VNT

The PPRV was diluted to 102.0 TCID50/0.1 mL with serum-free cell culture medium (MEM); the serum from the test group, positive control group and negative group, were Inactivated in a water bath at 56°C for 30 min. The non-immunized goat serum and negative serum were diluted 1:2, 1:4, 1:8 with MEM. The immunized goat serum and positive control serum were first diluted 1:4, and then make two-fold serial dilutions to 1:32. Each dilution of the serum was separately added to 5 wells of 96-well plate (0.1 mL/well). A virus regression experiment was also established. The virus working solution with 100 TCID50/0.1 mL was diluted with MEM to 10 TCID50/0.1 mL, 1 TCID50/0.1 mL, and 0.1 TCID50/0.1 mL. Each gradient virus solution was inoculated into 5 wells (0.1 mL/well). 0.1 mL of 100 TCID50/0.1 ml of virus working solution was added to all serum wells, 0.2 mL of MEM was added to the cell control well, 0.1 mL of MEM was added to the virus control well, and the mixture was allowed to incubate at 37°C for 1 hour. Next, add 0.1 mL of Vero cell suspension with a concentration of 200,000 to 300,000/mL to all wells and incubate for 6 days at 37°C in a 5% CO2 incubator. The plates were checked periodically using an inverted microscope and observation of a cytopathic effect (CPE) meant that the PPRV had not been neutralized by the serum dilution. Serum was considered positive for PPRV antibodies if the neutralizing dilution was greater than or equal to 1:10.

### Blocking Test

The blocking test (S5b Fig) was performed as previously reported with minor modification ^34^. First, the lyophilized peptides were dissolved with 30% acetonitrile solution (v/v, in pure water) to 1 mg/mL and further diluted to 0.5mg/mL with serum-dilution buffer; the proteins of N, F and H were diluted to 0.2 mg/mL with serum-dilution buffer. The serum T3-2 (18 dpv) diluted 1:100 with dilution buffer was mixed with equal volume of the diluted peptide or protein solution and incubated on a shaker for 30 min at room temperature (r.t.). 100 μL of the mixed solution was then transferred to the Microarray-#2 and processed as described in Serum screening with microarrays.

### Denaturing treatment

100 μL of 8M Urea or 8M urea & 20mM DTT in combination was added to the microarray-#2 to incubate for 30min at r.t. (S5a Fig). Then the microarray was rinsed for 3 times with washing buffer and incubated with specific serum samples, followed by steps as described in the blocking test.

### Field samples tested with Microarray-#2 (verification/validation groups

Large scale field samples were test with Microarray-#2. 100 μL of serum sample diluted 1 : 50 were incubated for 30 min (500 rpm, 37°C), rinsed for 3 times with washing buffer and incubated with of Rabbit anti-Goat IgG-HRP diluted 1:20000 in Peroxidase Conjugate Stabilizer/Diluent for another 30 min (500 rpm, 37°C), followed by the same washing step. 70 μL 3,3’,5,5’-tetramethylbenzidine (TMB) (Thermo scientific) was added onto the microarray and stands for 5 min to produce insoluble dark blue precipitate, followed by the wash with pure water and finally imaged by a scanner (ABT-XS01 imaging system, Suzhou Epitope, China) for chromogenic substrate. A positive control serum (PCS) responsive to all ECSPs and proteins and a negative control (NCS) serum were included in the test and the signal of individual ECSP or protein acquired by Matlab was further converted to the ratio of sample to positive control (S/P) using the formula below: S/P = (Signal (sample) – Signal (NCS)) / (Signal (PCS) – Signal (NCS)). All the Tests involving wild virus infected sera were performed in biosafety level-3 laboratory in Lanzhou Veterinary Research Institute, Chinese Academy of Agricultural Sciences.

### Statistical Analysis

The R package “pheatmap” was used for the cluster analysis and heat map performing. Spearman's rank correlation coefficient was calculated by R package to assess the repeatability of two experiments. Violin plot is performed using Graphpad Prism software and T tests were used to analyze the significant difference. Venn diagrams were drawn with tool of Venn Diagrams (http://bioinformatics.psb.ugent.be/webtools/Venn/). Receiver Operator Characteristic curve (ROCC) was performed with Graphpad Prism software.

## Acknowledgments

This study was supported by National Natural Science Foundation of China (No. 31602035) (to Q.X.) and the project of National Key R&D Program of China 2016YFD0500800 (to H.M.). We would like to thank Mrs. Lan Yang for the help with bioinformatics analysis.

## Author Contributions

H. M., X. C., and Q. X. conceived the study; H. X., H. L., J. P., M. S. and Y. C. performed all the experiments; H. X., H. L., Q. X. and H. M. analyzed the results; J. Y. analyzed the results related to the structure of all proteins; H. X., W. X., Q. X. and H. M. wrote the manuscript. All authors contributed to the revision and review of the manuscript.

## Supplemental files

S1 Text. The urgent demand for DIVA strategy cooperated with live-attenuated vaccines.

S2 Text. Detailed analysis of the remaining goats of group T1, T2 and T3 respectively.

S3 Text. Effect of denaturing agents on the structural proteins of PPRV.

S4 Text. The advantages and the remaining challenges of Microarray-#2.

S1 Fig. Photographs of real objects (Microarray-#1/-#2).

S2 Fig. The IgG serodynamics of three goats (T1) for early stage (0-21 dpv).

S3 Fig. The IgG serodynamics of three goats (T2) for the period of 0-44 dpv.

S4 Fig. The IgG serodynamics of T3-1 and T3-3 for the period of 0-228 dpv.

S5 Fig. Denaturation and blocking experiments.

S6 Fig. SDS-PAGE and Western Blot of N, H and F protein.

S7 Fig. ROC curves of four ECSPs and three proteins.

S8 Fig. Test results of field samples on individual ECSP or protein level.

S9 Fig. Venn diagrams of the positive results of C1 and negative results of C2&C3 based on ECSPs and Proteins.

S10 Fig. Using PPR live-attenuated vaccine to mimic the wild virus infection.

S1 Table. Examples of diseases caused by viruses: eradication related status.

S2 Table. A DIVA microarray strategy works for any vaccine.

