## Supplemental files for "Epitope containing short peptides capture distinct IgG serodynamics that enable DIVA for live-attenuated vaccines"

**This file includes:**

S1 to S4 Text

S1 to S10 Fig

S1 to S2 Table

**S1 Text. The urgent demand for DIVA strategy cooperated with live-attenuated vaccines.**

Outbreaks of infectious diseases caused by viruses, such as Pseudorabies (PR) Foot-and-mouth disease (FMD), and Peste des petits ruminants (PPR), led economic losses reaching billions of dollars (S1 Table).

Both PR and FMD were eliminated in several countries via large-scale eradication programs, which first control the disease through vaccination then ultimately remove the transmission of field strain of viruses through DIVA. PR has a DIVA-compatible vaccine that lacks the gE protein (i.e., the negative marker, NM) and a companion diagnostic test that uses the gB protein as the marker (M) and gE as the NM. FMD has a DIVA-compatible vaccine that has non-structural proteins (NSPs) as NM and structural proteins (SPs) as the marker (M). For vaccinated herds, results from its companion diagnostic test are given as (anti-NM IgG, anti-M IgG), (-, +) and (+, +) indicated vaccinated and infected samples, respectively [6, 7].

Live-attenuated vaccines elicit not only the best protection, but also represent the most time-saving method for vaccine development [31]. Although they do come with a risk for toxic resilience, live-attenuated vaccines have many advantages, including low dosages, good immunogenicity, a long protection period, low cost, and convenient use. For example, there was an outbreak of PPR in 2007 in China [35], yet the use of live-attenuated vaccine (PPRV Nigeria 75/1 strain) was quickly put in use, and this was the key in preventing PPRV causing further devastation.

The Office International Des Epizooties highly recommended two methods for PPR diagnosis, especially for the evaluation of post vaccination immune status, namely a competitive ELISA (cELISA) and a virus neutralization test (VNT) [10]. Unfortunately, both are incompatible with DIVA. More than 10 years has passed since the first outbreak of PPR in China, and the live-attenuated vaccine is still in use. PPR causes >2 billion USD losses annually worldwide [36]. To realize the goal of global eradication of PPR by 2030 [37] and China's goal of national elimination by the end of 2020, a DIVA solution is urgently needed. Considering PPR is still prevalent in most of Africa, the Middle East, and South Asia, especially in countries surrounding China [38], a DIVA solution is also needed in the long run for monitoring, even after successfully eliminating PPR from China.

**S2 Text. Detailed analysis of the remaining goats of group T1, T2 and T3 respectively.**

The 9 goats were divided into three groups, groups T1 and T2 were immunized with a single PPR vaccine and group T3 was immunized with a combined PPR vaccine. They were followed by different monitoring protocols: 1) group T1 sampled sera daily for a period of 0-21 dpv; 2) T2 sampled sera at 3–4 day intervals for a period of 0-44 dpv and 3) T3 sampled at 0 dpv and at 3–4 day intervals for a period of 11–228 dpv (S2a, S3a and S4a Fig). All 9 goats showed a similar trend with minor discrepancies due to individual immune differences.

Sera from T1 group (sampled daily) provided detailed information on the initial development of the humoral immune response. Comparison of IgG serodynamic curves determined by cELISA and VNT demonstrated antibodies against N protein appeared slightly earlier than neutralization antibodies (7dpv vs. 8-10dpv) (S2d-e Fig), which were consistent with a previous report [17].

For goat T1-1, anti-N49 IgGs first appeared at 9 dpv while anti-F10 IgGs first appeared at 17 dpv. For goat T1-3, anti-N49 IgGs showed 3 times higher signal intensity than anti-F10 IgGs. The IgG serodynamics also showed significant individual immune differences among the 3 goats. For the first appearance of anti-N49 IgGs, goat T1-1 was 2-3 days earlier than the other two goats. For the first appearance of anti-F10 IgGs, however, goat T1-1 was 5-7 days later than the other two goats. cELISA produced relatively uniform results: all three goats became positive at 7 dpv and were strongly positive after 9 dpv. VNT tests indicated T1-1, T1-2, and T1-3 showing exhibiting positive readouts at 10, 8, and 9 dpv, respectively. They all became strong and steadily positive after 14dpv.

Compared with the T1 group, sera from the T2 group not only prolonged the sampling periods (21 dpv vs. 44 dpv) but a longer interval (daily vs. every 3-4 days). The results derived, however, were similar to that from T1 (S3 Fig). Thus, T3 was designed according to T2's protocol.

Sera from the T3 group were used to monitor long-term immune responses. Results from T3-1 and T3-3 indicated the similar curves for anti-N49 and anti-F10 IgGs with slight differences in the peak times. ECSPs N45-N50 and F9-F11 showed strong response in all 9 goats during the 10-60 dpv period, peaking around 20 dpv (S2-4 Fig).

If one assumes all IgGs have the same affinity, the signal difference reflects the concentration difference. It is obvious that IgGs elicited by the live-attenuated vaccine at 18 dpv consist of 10 times more anti-protein IgGs than anti-ECSP IgGs (Fig 2e).

**S3 Text. Effect of denaturing agents on the structural proteins of PPRV.**

For the denaturation experiments, immobilized ECSPs (N47/49/50, F10) or proteins on Microarray-#2 were first denatured using 8M urea or combined 8M urea plus 20mM DTT. Assuming the conformational structures were destroyed due to the treatments, the microarray were reacted with serum T3-2 (18 dpv), which was collected at the early stage of immunization. See Fig 2e-f for data.

Although no structures of PPRV proteins are available so far, the crystal structures of the other morbilliviruse MeV proteins have been extensively studied [39]. Taking the MeV structures as the template, we could build the hypothetical structural models of PPRV proteins considering that their protein sequence identities are up to 50% or more (Structural models for PPRV proteins were not shown in this paper).

For example, the structure of PPRV H is also a tetramer that comprises two dimers and the binding surface of the two dimers are mainly relied on hydrophobic interactions as MeV H [40]. It is widely accepted that urea is able to attack hydrophobic residues of protein. When treating PPRV H protein with urea, it could first break the hydrophobic interface of the tetramer by enhancing the solubility of interacting molecules; subsequently influencing the structures of conformational epitopes that interacted with restricted IgGs of the T3-2 (18 dpv) serum.

Consistent with previous work, F protein of PPRV contains 16 cysteines and at least five pairs of disulfide bonds are really essential for its structure maintenance, which is also the case for MeV F. Based on this information, it seems easier to destroy the structure of F protein and make it denatured by using DTT rather than urea, which was in accord with above data, further implying that the IgGs in this immune stage apparently have few linear epitopes.

**S4 Text. The advantages and the remaining challenges of Microarray-#2.**

For individual ECSPs or proteins, the result was positive if its S/P was larger than the cutoff; for the ECSPs combination (i.e. N47+N49+N50+F10), the result was positive if any ECSP was positive; for Microarray-#2, the result was positive if any ECSP was positive or any two proteins were positive. Although the ROC curves showed the diagnostic performance of individual ECSPs did not singly meet typical thresholds (both of sensitivity and specificity should be more than 90%), it is possible to use a combination of four ECSPs to fulfill this industrial standard. Although the specificity of all four ECSPs were apparently perfect, the sensitivity of individual ECSP was far below the 90% standard needed for a diagnostic biomarker. This drawback was remedied by the use of four ECSPs jointly.

Since this project lasted for more than 2 years from the first serum sample was collected, most of the donors (goats) no longer exist. We were thus unable to do more tests. A large scale validation test for the DIVA diagnostic microarray has been initiated.

For those DIVA-compatible vaccines, this DIVA microarray strategy could be used to replace the accompanied diagnostic kits, mainly ELISA, that detect negative markers to indicate infection.

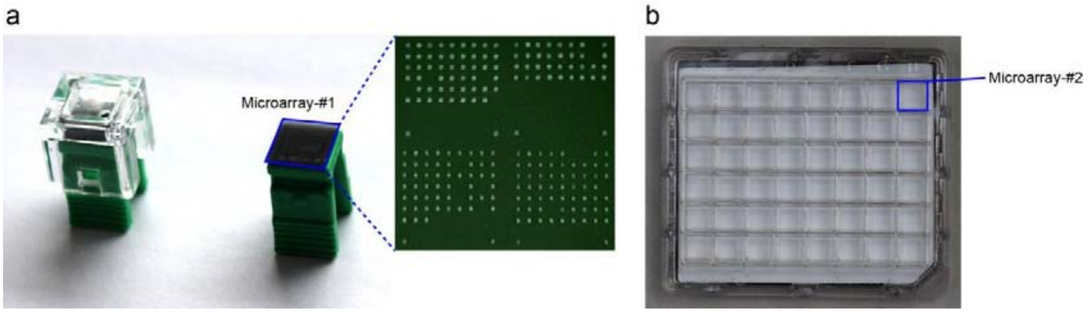


**S1 Fig. Photographs of real objects (Microarray-#1/-#2).** (a) Microarray-#1 was designed for individual serum screening use. A nano-membrane printed with Microarray-#1 was first assembled to the plastic support (right), and then covered with a plastic lid to form the reaction chamber (left); (b) Microarray-#2 was designed for large scale screening use, each plate comes with 48 Microarray-#2.

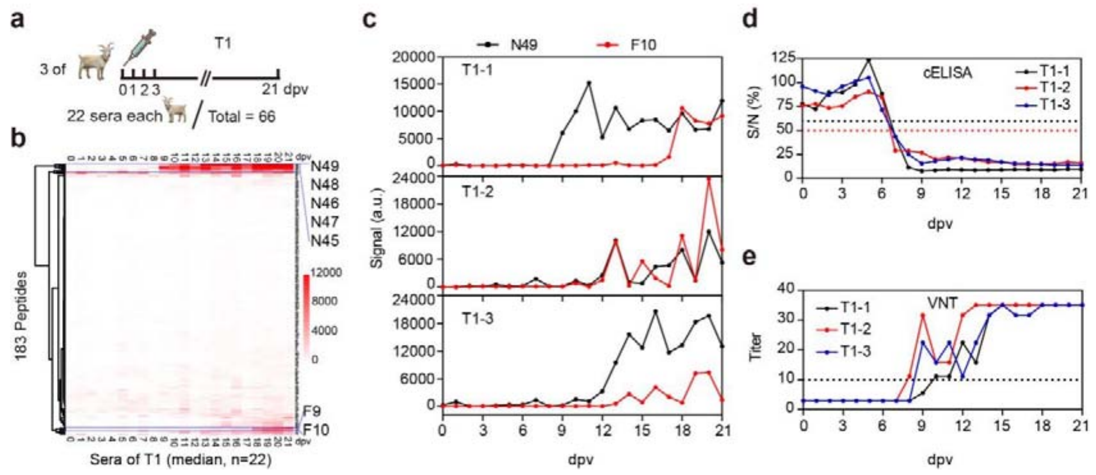


**S2 Fig. The IgG serodynamics of three goats (T1) for early stage (0-21 dpv).** (a) 3 goats were immunized with a single shot of PPR vaccine and sampled for sera daily. 22 sera samples were collected for each goat; (b) All 66 sera were screened using Microarray-#1. Median value calculated from three sera at the same dpv was used to construct a heat-map, from which two groups of ECSPs were seen to increase in signal intensity over time, namely F9-10 and N45-N49; (c) Individual goats showed different seroconversion dynamics for ECSPs; Results from (d) cELISA and (e) VNT tests.

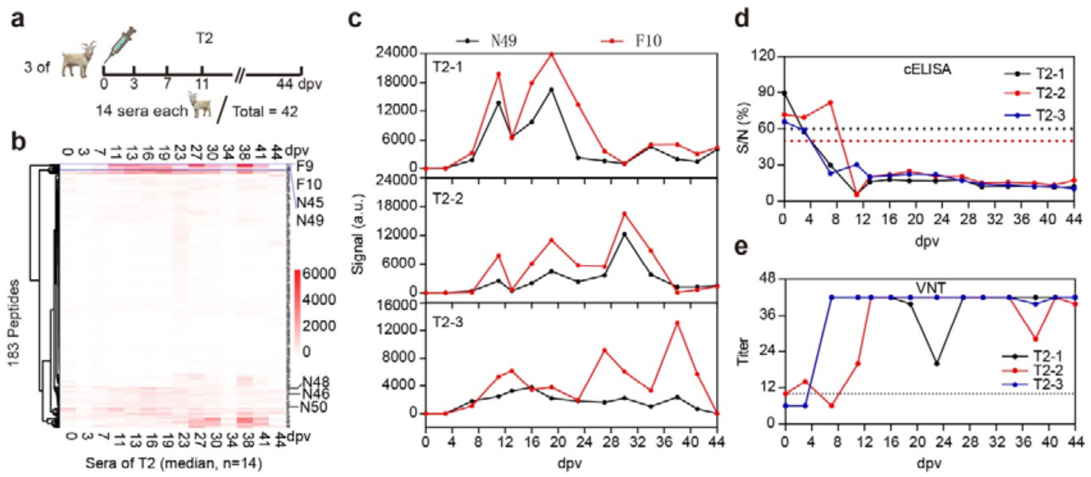


**S3 Fig. The IgG serodynamics of three goats (T2) for the period of 0-44 dpv.** (a) 3 goats were immunized with a single shot of PPRV vaccine and sampled every 3-4 days. Each goat produced 14 sera; (b) All 42 sera were screened against Microarray-#1, resulted in a heat-map, from which the same ECSPs were seen to first increase then decline over time, namely F9-10 and N45-N49; (c) T2-1 was significantly different from the other two, it became IgG positive at 7 dpv and peaked at 19 dpv, abbreviated as (7, 19). T2-2 and T2-3 showed (11, 30) and (7, 40); (d) cELISA produced relatively uniform results: T2-1 and T2-3 became positive at 7 dpv and T2-2 at 11 dpv. They all became strongly and steady positive after 13 dpv; (e) VNT tests indicated the same trends as the cELISA.

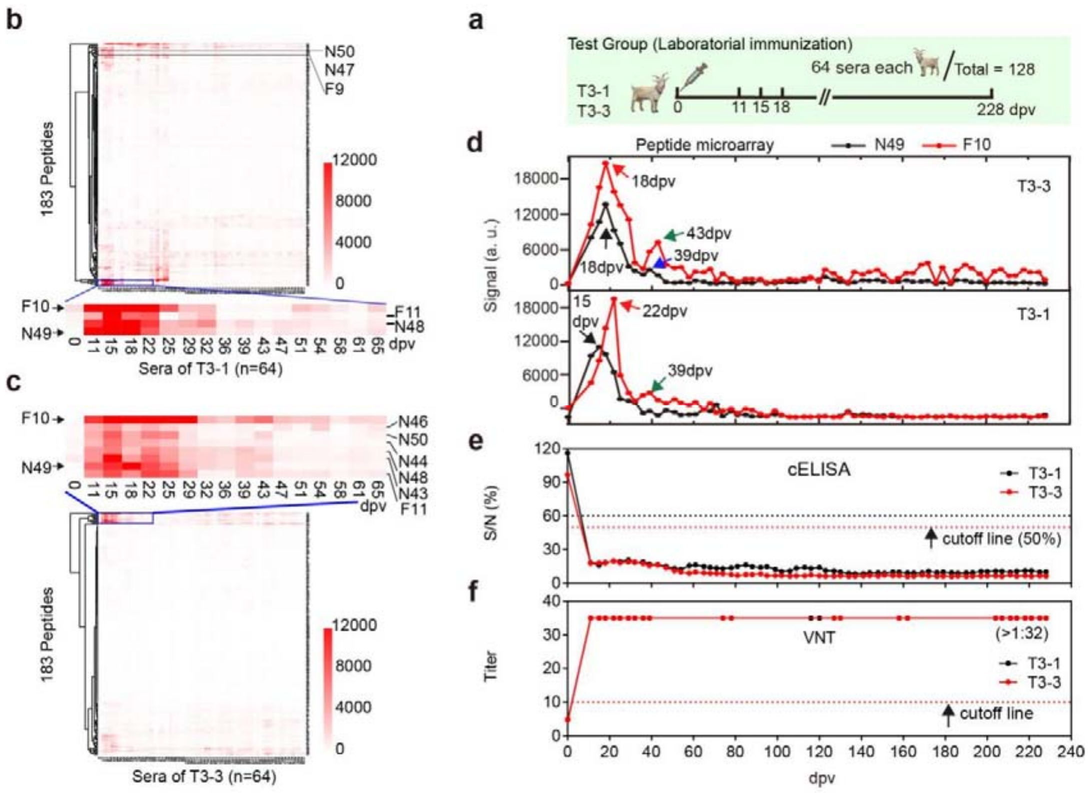


**S4 Fig. The IgG serodynamics of T3-1 and T3-3 for the period of 0-228 dpv.** (a) They were immunized with a combined PPR vaccine and sampled every 3-4 days. Each goat produced 64 sera. All 128 sera were screened against the same Microarray-#1, resulted in a heat-map for (b) T3-1 and (c) T3-3; (d) The IgG serodynamics detected based on ECSPs F9-10 and N45-N49. The trend as defined by the first appearance of a positive sign; peaks and valleys were similar: anti-F10 IgGs peaked at 15 and 18 dpv, anti-N49 IgGs peaked at 22 and 18 dpv for T3-1 and T3-3 respectively; (e) cELISA and (f) VNT tests both indicated complete seroconversion at 11 dpv.

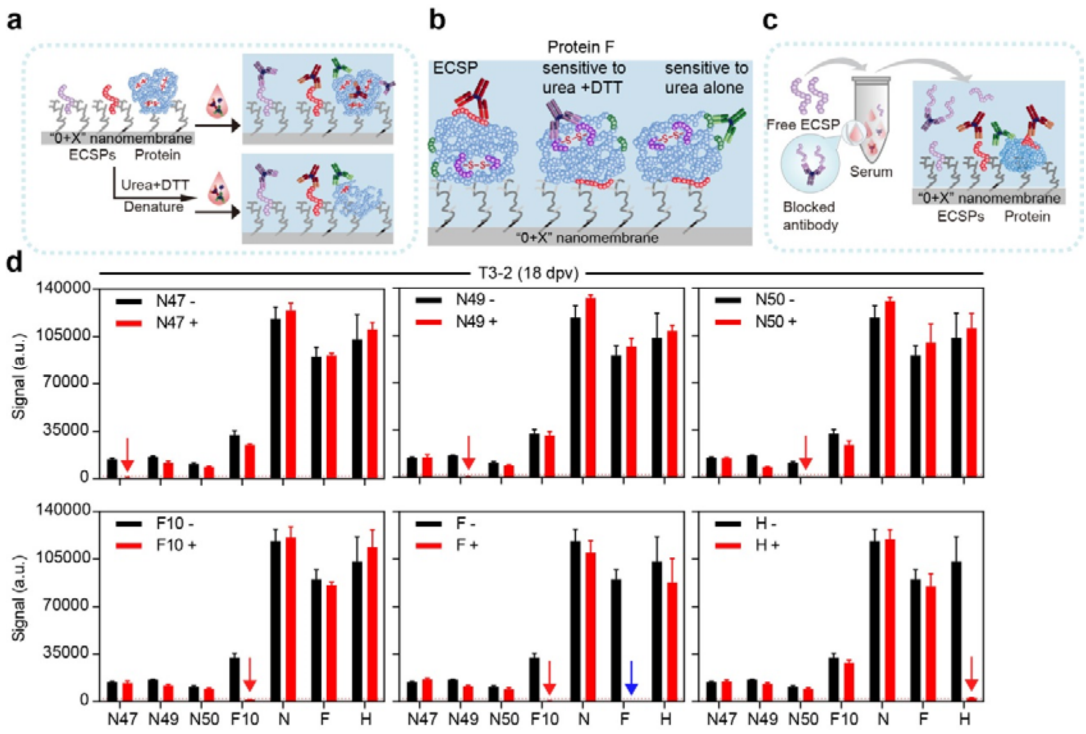


**S5 Fig. Denaturation and blocking experiments.** (a) A flowchart of denaturation experiments (see Fig 2e-f for data); (b) A flowchart of blocking experiments; (c) results of blocking experiments. Free ECSP or protein was individually doped to serum before screening against Microarray-#2; (d) Without denaturation treatment (black column), T3-2 showed considerable responses to ECSPs and especially to N, F, and H proteins.

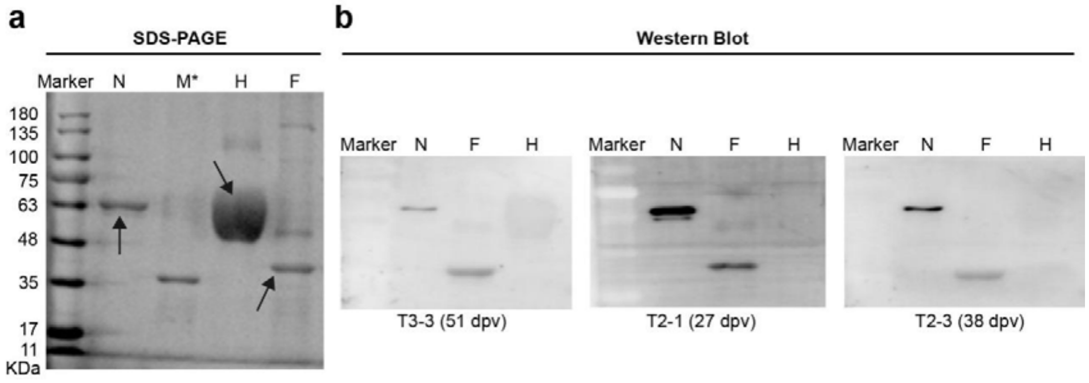


**S6 Fig. SDS-PAGE and Western Blot of N, H and F protein.** (a) Polyacrylamide gel electrophoresis (SDS-PAGE) analysis of Protein N, H, and F using Coomassie Blue staining. The molecular weight (MW) of Protein N, H, and F is about 60kDa, 66kDa, and 52kDa respectively, as pointed by arrow. Protein F is present in both monomeric and oligomeric forms. (b) Western blot analysis of the antigenicity of Protein N, H, and F. Serum samples collected at different dpv showed good reactivity to Protein N and F but showed little reactivity to Protein H, implying that Protein N and F could have the corresponding linear epitopes, while Protein H did not have rich linear epitopes, which were consistent with previous data.

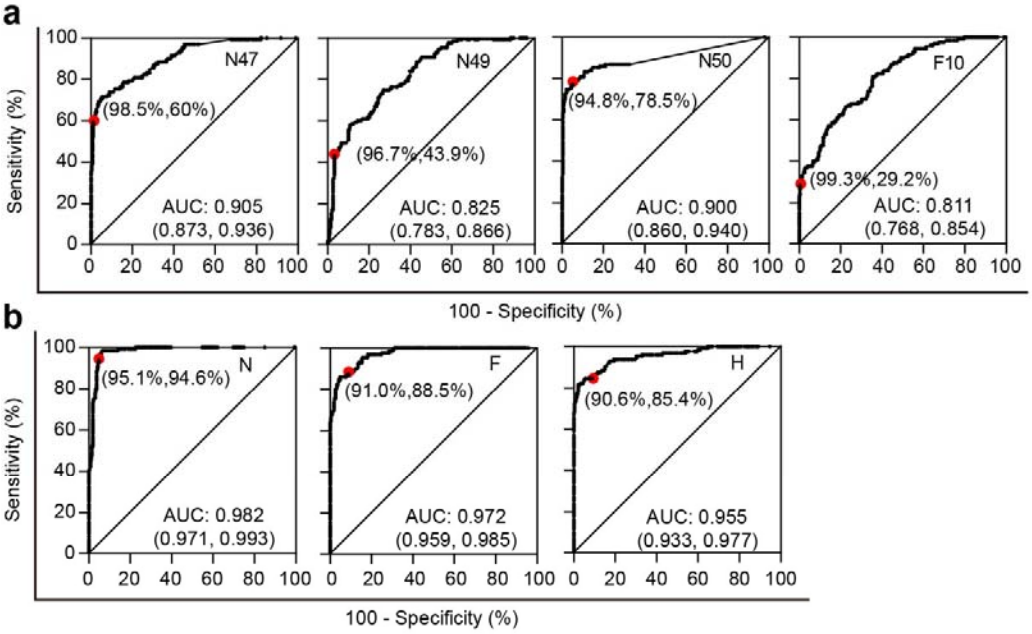


c

| ECSP/ Protein | S/P_cutoff | Sensitivity% | 95% CI | Specificity% | 95% CI |
|---|---|---|---|---|---|
| N47 | 0.4 | 60 | 51.05% to 68.49% | 98.5 | 96.21% to 99.59% |
| N49 | 0.45 | 43.85 | 35.16% to 52.82% | 96.63 | 93.70% to 98.45% |
| N50 | 0.15 | 78.46 | 70.40% to 85.19% | 94.76 | 91.36% to 97.10% |
| F10 | 0.7 | 29.23 | 21.59% to 37.85% | 99.25 | 97.32% to 99.91% |
| N | 0.3 | 94.62 | 89.22% to 97.81% | 95.13 | 91.82% to 97.38% |
| F | 0.2 | 88.46 | 81.68% to 93.40% | 91.01 | 86.92% to 94.16% |
| H | 0.12 | 85.38 | 78.12% to 90.97% | 90.64 | 86.49% to 93.85% |

**S7 Fig. ROC curves of four ECSPs and three proteins.** (a) ROC curves of N47, N49, N50, and F10 obtained from 267 negative sera and 130 vaccinated sera at 29 dpv; (b) ROC curves of N, F, and H proteins. The red points represented the specificity and sensitivity respectively at the corresponding S/F cutoff values. (c) Values for the red points of curves in (a) and (b).

**a** C1 (Negative, n=267)

**b** C2&C3 (n=130) ~29 dpv → C4 (n=8) 480 dpv → C5 (n=17) 930 dpv

**c** C6 (infected, n=25)

| | | C1 Neg. | C1 Pos. | C2&C3 Neg. | C2&C3 Pos. | C4 Neg. | C4 Pos. | C5 Neg. | C5 Pos. | C6 Neg. | C6 Pos. |
|---|---|---|---|---|---|---|---|---|---|---|---|
| ECSP | N47 | 264 | 3 | 52 | 78 | 6 | 2 | 9 | 8 | 12 | 13 |
| ECSP | N49 | 258 | 9 | 73 | 57 | 8 | 0 | 15 | 2 | 15 | 10 |
| ECSP | N50 | 253 | 14 | 28 | 102 | 8 | 0 | 11 | 6 | 16 | 9 |
| ECSP | F10 | 266 | 1 | 92 | 38 | 8 | 0 | 15 | 2 | 23 | 2 |
| | ECSPs | 242 (90.6%) | 25 | 13 | 117 (90%) | 6 | 2 | 6 | 11 | 7 | 18 |
| Protein | N | 253 | 14 | 7 | 123 | 0 | 8 | 1 | 16 | 0 | 25 |
| Protein | F | 243 | 24 | 16 | 114 | 0 | 8 | 0 | 17 | 3 | 22 |
| Protein | H | 241 | 26 | 19 | 111 | 0 | 8 | 0 | 17 | 0 | 25 |

**d**

| | 1 | 2 |
|---|---|---|
| N47 | 1.265 | 0.455 |
| N49 | -0.095 | -0.241 |
| N50 | -0.169 | -0.263 |
| F10 | -0.483 | -0.035 |
| N | 0.855 | 0.642 |
| F | 1.050 | 0.678 |
| H | 1.047 | 0.967 |

| VNT | cELISA | Microarray-#2 Protein | Microarray-#2 ECSPs | NO. | Microarray-#1 | Total | F | H | M | N |
|---|---|---|---|---|---|---|---|---|---|---|
| + | + | + | + | 1 | + | 33 | 4 (1) | 10 (3) | 1 | 18 (9) |
| + | + | + | + | 2 | + | 38 | 11 (10) | 9 (5) | 9 (7) | 9 (9) |

**e**

| | 1 | 2 | 3 | 4 | 5 | 6 | 7 | 8 | 9 | 10 | 11 |
|---|---|---|---|---|---|---|---|---|---|---|---|
| N47 | 1.19 | 0.94 | -0.09 | -0.40 | 0.15 | 0.48 | -0.05 | 0.32 | 0.48 | 0.78 | 0.25 |
| N49 | -0.63 | 0.00 | -0.15 | -0.25 | -0.27 | 0.29 | -0.47 | -0.41 | -0.46 | 0.05 | -0.63 |
| N50 | -0.15 | -0.15 | 0.73 | 0.03 | 0.14 | -0.15 | 0.01 | -0.02 | 0.03 | 0.83 | -0.15 |
| F10 | -0.25 | -0.41 | 0.22 | -0.43 | -0.25 | -0.46 | -0.93 | -0.60 | -0.71 | 0.09 | -0.22 |
| N | 0.29 | 0.42 | 0.90 | 0.46 | 0.43 | 0.53 | 0.50 | 0.41 | 0.15 | 0.64 | 0.41 |
| F | 0.50 | 0.83 | 0.92 | 0.82 | 0.79 | 0.79 | 1.00 | 0.91 | 0.24 | 0.82 | 1.05 |
| H | 0.41 | 0.89 | 0.95 | 0.80 | 0.74 | 0.73 | 0.93 | 0.76 | 0.65 | 0.81 | 1.12 |

| VNT | cELISA | Microarray-#2 Protein | Microarray-#2 ECSPs | NO. | Microarray-#1 | Total | F | H | M | N |
|---|---|---|---|---|---|---|---|---|---|---|
| + | + | + | + | 1 | + | 64 | 27 (4) | 15 (2) | 6 (3) | 16 (2) |
| + | + | + | + | 2 | + | 46 | 14 (4) | 9 (3) | 9 (1) | 14 (4) |
| + | + | + | + | 3 | + | 18 | 7 (3) | 3 (1) | 2 (1) | 6 (2) |
| + | + | + | + | 4 | + | 15 | 2 | 4 | 5 | 4 (2) |
| + | + | + | + | 5 | + | 11 | 1 | 3 | 3 (2) | 4 (1) |
| + | + | + | + | 6 | − | 5 | 3 | 0 | 0 | 2 |
| + | + | + | + | 7 | − | 4 | 1 | 3 | 0 | 0 |
| + | + | + | + | 8 | − | 2 | 1 | 0 | 0 | 1 |
| + | + | + | + | 9 | − | 1 | 0 | 0 | 0 | 1 |
| + | + | + | + | 10 | − | 0 | 0 | 0 | 0 | 0 |
| + | + | + | + | 11 | − | 0 | 0 | 0 | 0 | 0 |

**f**

| | 1 | 2 | 3 | 4 | 5 | 6 | 7 |
|---|---|---|---|---|---|---|---|
| N47 | -0.40 | -0.10 | -0.24 | -0.32 | -0.30 | -0.24 | -0.30 |
| N49 | -0.42 | -0.56 | -0.48 | -0.44 | -0.62 | -0.12 | -0.62 |
| N50 | -0.26 | -0.26 | -0.17 | -0.15 | -0.26 | -0.17 | -0.17 |
| F10 | -0.25 | -0.43 | -0.50 | -0.48 | -0.67 | -0.50 | -0.63 |
| N | 0.46 | 0.48 | 0.59 | 0.61 | 0.49 | 0.75 | 0.33 |
| F | 0.63 | 0.89 | 0.54 | 0.67 | -0.37 | 0.25 | 0.10 |
| H | 0.68 | 0.89 | 0.84 | 0.92 | 0.25 | 0.40 | 0.09 |

| VNT | cELISA | Microarray-#2 Protein | Microarray-#2 ECSPs | NO. | Microarray-#1 | Total | F | H | M | N |
|---|---|---|---|---|---|---|---|---|---|---|
| + | + | + | − | 1 | + | 28 | 7 (1) | 5 (2) | 6 (2) | 10 (1) |
| + | + | + | − | 2 | + | 30 | 0 | 0 | 14 (4) | 16 (5) |
| + | + | + | − | 3 | + | 47 | 10 (1) | 0 | 13 (4) | 24 (9) |
| + | + | + | − | 4 | + | 41 | 7 (3) | 0 | 8 (2) | 26 (9) |
| + | + | + | − | 5 | + | 24 | 3 (1) | 0 | 7 (3) | 14 (4) |
| + | + | + | − | 6 | + | 39 | 5 (2) | 0 | 9 (1) | 25 (8) |
| + | + | + | − | 7 | + | 22 | 2 | 0 | 8 (4) | 12 (3) |

**S8 Fig. Test results of field samples on individual ECSP or protein level.** (a) C1, the 267 negative sera, (b) Left: 130 positive sera (C2 and C3 combined), Middle: C4, 8 positive sera at 480 dpv, Right: C5, 17 positive sera at 930 dpv, (c) C6, 25 positive sera due to wild virus infection, (d) Detailed results of 2 sera of C4 with positive ECSPs response, (e) Detailed results of 11 sera of C5 with positive ECSPs response, (f) Detailed results of 7 sera of C6 with negative ECSPs response.

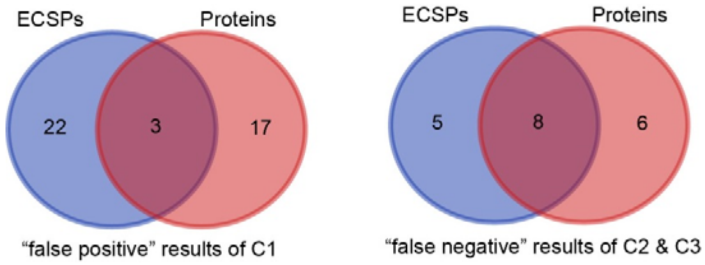


**S9 Fig. Venn diagrams of the positive results of C1 and negative results of C2&C3 based on ECSPs and Proteins.** 0, 3, and 0 out of the 25 sera of C1 were shown to be negative by N, F, and H proteins respectively, implying N and H proteins are more appropriate than F protein for the diagnosis of wild infectious cases.

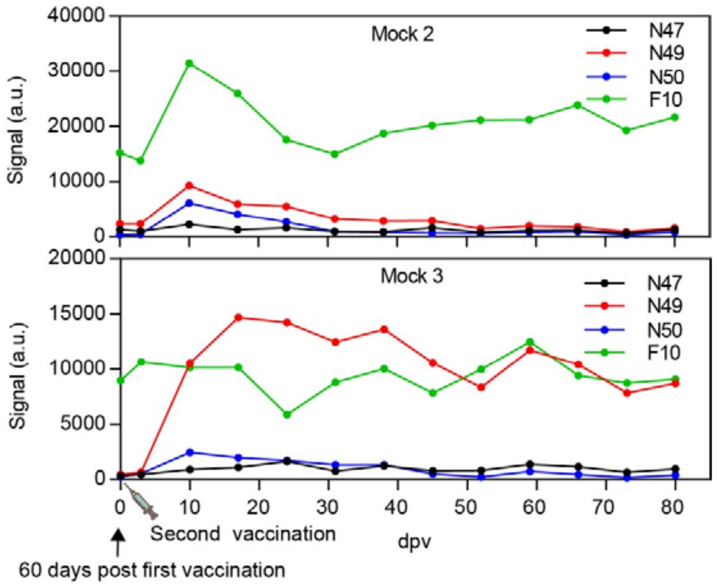


**S10 Fig. Using PPR live-attenuated vaccine to mimic the wild virus infection.** The spiked ECSPs response clearly indicated "infection". The difference we saw in Mock 2 and Mock 3 was due to individual immune differences.

**S1 Table. Examples of diseases caused by viruses: eradication related status.**

| Disease | Vaccine (DIVA) | DIVA test | | Eradication status | | Outbreak | No. of slaughtered animals | Losses (USD) |
|---|---|---|---|---|---|---|---|---|
| | | M | NM[a] | Succeeded | Epidemic | | | |
| PPR | Live attenuated ( - ) | | N/A | N/A | ~70 countries | 2007, China | N/A | 2.1 billion/Y (Worldwide) |
| FMD | Inactivated (without NSP) ( + ) | SPs | NSPs[b] | USA, Canada, Australia, et al | China, Egypt, Vietnam, et al | 2001, England | 6.5 million | 13.8 billion |
| PR | Inactivated (gene deletion) ( + ) | gB | gE | USA, Germany, Denmark, et al | China, India, Vietnam,et al | 1990, USA | N/A | 0.6 billion |

[a]NM: Negative marker ; [b]NSPs: Non-structural proteins.

**S2 Table. A DIVA microarray strategy works for any vaccine.**

| Vaccine type | A DIVA vaccine | Example (Disease) | Traditional DIVA test (Negative maker) | A DIVA microarray |
|---|---|---|---|---|
| Live attenuated vaccine | NO | PPR | N/A | A distinct IgG serodynamic |
| Inactivated whole virus vaccine | YES | FMD/AI | NSP/Different strain | YES |
| Subunit vaccine | YES | CSF | Erns | YES |
| Gene reverse engineering | YES | PR | gE | YES |
